# Optimization of Neuroprosthetic Vision via End-to-end Deep Reinforcement Learning

**DOI:** 10.1101/2022.02.25.482017

**Authors:** Burcu Küçükoğlu, Bodo Rueckauer, Nasir Ahmad, Jaap de Ruyter van Steveninck, Umut Güçlü, Marcel van Gerven

## Abstract

Visual neuroprostheses are a promising approach to restore basic sight in visually impaired people. A major challenge is to condense the sensory information contained in a complex environment into meaningful stimulation patterns at low spatial and temporal resolution. Previous approaches considered task-agnostic feature extractors such as edge detectors or semantic segmentation, which are likely suboptimal for specific tasks in complex dynamic environments. As an alternative approach, we propose to optimize stimulation patterns by end-to-end training of a feature extractor using deep reinforcement learning agents in virtual environments. We present a task-oriented evaluation framework to compare different stimulus generation mechanisms, such as static edge-based and adaptive end-to-end approaches like the one introduced here. Our experiments in Atari games show that stimulation patterns obtained via task-dependent end-to-end optimized reinforcement learning result in equivalent or improved performance compared to fixed feature extractors on high difficulty levels. These findings signify the relevance of adaptive reinforcement learning for neuroprosthetic vision in complex environments.

## 1 Introduction

Globally, approximately 50 million people suffer from blindness, and 250 million are visually impaired. Many may benefit from neuroprostheses that bypass the damaged area in the visual pathway and directly stimulate a small subset of cells in the retina, axons in the optic nerve, or neurons in the visual cortex [1, 2]. Through artificially induced electrical stimulation, prosthetic vision enables perception of coarse light dots called *phosphenes* [3]. Spatial coherence is maintained due to the retinotopic organization of the primary visual cortex [4]. A challenge with phosphene vision through electric stimulation is to extract a compressed yet informative stream of sensory information from the complex surroundings. Sparse stimulation patterns are desirable to minimize the energy consumption of the neuroprosthesis and to maintain stimulation intensities that are sustainable by the neural tissue in the long term [5, 6]. Previous research [7] has focused on direct mechanistic image translation from camera footage to stimulation patterns, often using a single kind of feature, such as edges, segmentation maps, or keypoint detection (see e.g. [8, 9, 10]). In these studies, each type of feature appears well suited for a particular scenario [11] such as navigation, object detection, or emotion recognition. Our hypothesis is that realistic environments require a more tailored approach in which the stimulation patterns are optimized for the task at hand.

According to surveys among visually impaired people, a major concern is the restriction of their mobility in unfamiliar environments [1]. Deep reinforcement learning (RL) has been very successful in training virtual agents to perform multi-step goal-directed behavior in complex dynamic environments. Inspired by these advances, we make two contributions. First, we developed a framework to evaluate stimulus generation mechanisms in task-oriented simulated environments using Reinforcement Learning (RL). Second, we developed an optimization method for Task Optimized Phosphene Vision (TOPhos), where optimal stimulation patterns are learnt directly from interactions with the environment.

The optimization method consists of a deep neural network that encodes stimulation patterns, referred to as the encoder. A phosphene simulator transforms the stimulation patterns into simulated phosphene vision, a representation of what blind people might see in response to electrical stimulation. A virtual agent receives this phosphene vision as sensory input and uses it to solve tasks involving navigation, obstacle avoidance, object approaching, and high-level objectives such as replenishing a certain resource. The reward signals obtained while interacting with the environment are used to optimize the stimulation patterns (task objective). In addition to the RL agent, a decoder network uses the simulated phosphene vision to reconstruct the original image. The difference between original and reconstructed image informs about the phosphene quality and is used as an auxiliary learning signal in addition to the RL loss (reconstruction objective). The two loss terms are used to update all components of the architecture in an end-to-end fashion, simultaneously optimizing the task-dependent reward from the environment and the phosphene quality from the reconstruction.

We expect that the use of both task and reconstruction objectives to learn phosphene vision encoders will lead to phosphene patterns that are better adapted to dynamic multi-stage mobility scenarios faced by agents in the real world. As a proof of concept, we trained agents in visually simple but popular test beds of RL methods, namely Atari games, which combine navigation, approach and avoidance, object detection, and managing multiple simultaneous goals. To evaluate the efficacy of the proposed optimization framework, we compared its performance to baselines from (1) an RL agent with normal vision, (2) a blind RL agent, (3) an RL agent with fixed phosphene vision generated by edge-detection, (4) an RL agent whose phosphene vision is optimized without a task objective, and (5) an RL agent whose phosphene vision is optimized without a reconstruction objective.

### 1.1 Related Work

As most visual neuroprostheses are still in the development phase, the effect of stimulation patterns on functional gains is largely investigated using simulation studies. Previous implementations of simulated prosthetic vision use preprocessing techniques such as pixelization [8], feature extraction techniques to highlight the most relevant parts of the image based on saliency cues [9], environmental structure [10], or edge filtering [12, 13, 14]. Study [15] focused on implementation of more realistic phosphene maps, while [16] employed facial landmark detection for emotion recognition. In previous work [17], an autoencoder was trained where the encoder generated stimulation patterns for simulated phosphene vision and the decoder used the phosphenes to produce a reconstruction of the sensory input. The objective function used during optimization consisted of a reconstruction loss term between original and reconstructed sensory input, and a sparsity loss term to ensure economic electrode activation.

These approaches act on single static images and focus either on correct image reconstruction from phosphenes or design feature extractors to suit a particular task. In contrast, our work proposes an optimization procedure to learn a feature extractor that solves a given task optimally. Another difference is that whereas these previous approaches considered image-wise transformations, our RL-based architecture takes into account the task dynamics over extended periods of time.

In dynamic settings requiring sequential control of actions, RL has been applied to find optimal deep brain stimulation patterns for treating Parkinson’s disease [18] and epilepsy [19]. In the case of vision prostheses, we face a significantly higher dimensionality of the input and output space, as well as the lack of a well-defined target state. The present work is a step towards closing this gap. Deep reinforcement learning (DRL), which combines deep learning with reinforcement learning, provides benefits in addressing the issue of high dimensionality with the use of neural networks for learning latent representations of the input space. However, learning the representation through typically sparse task-specific rewards can lead to overfitting that prevents the model from generalizing to other tasks [20]. One way to address this issue is to include an auxiliary output unrelated to the main task [21]. Such an auxiliary output, for example from a secondary depth prediction task, has been shown to improve navigation of DRL agents in complex environments [22] as well as learning saliency maps for prosthetic vision [23]. Previous work used an image reconstruction task from phosphene patterns to optimize prosthetic vision [17]. Our approach employs this image reconstruction as an auxiliary task next to an RL agent introduced here for adaptive task-dependent representation learning.

## 2 Methods

### 2.1 RL as a Task-oriented Phosphene Vision Evaluation Framework

Here we propose a new method for evaluating the percept quality which a visual neuroprosthesis could provide. In this method, mobile virtual agents with phosphene vision act on their simulated environments to achieve some task that requires sequential decision making. Unlike phosphene vision evaluation methods that focus merely on image-based details in static test-beds, this task-oriented RL framework for evaluation of phosphene vision enables investigation of the functional gains different stimulus generation approaches could provide.

The framework (Fig.1) can be generally described by four elements: (1) a simulated virtual environment that provides the visual setting and the task, (2) a stimulus generation method that converts visual inputs from the environment scene to a stimulation pattern for inducing phosphenes through the neuroprosthesis, (3) a phosphene simulator that captures the perceptual effect of the stimulation pattern to compose the vision of the agent, and (4) a reinforcement learning agent that learns to act towards task goals based on its phosphene vision and sequential interaction with the environment. on mobility and goal-directed behavior.

**Figure 1:**
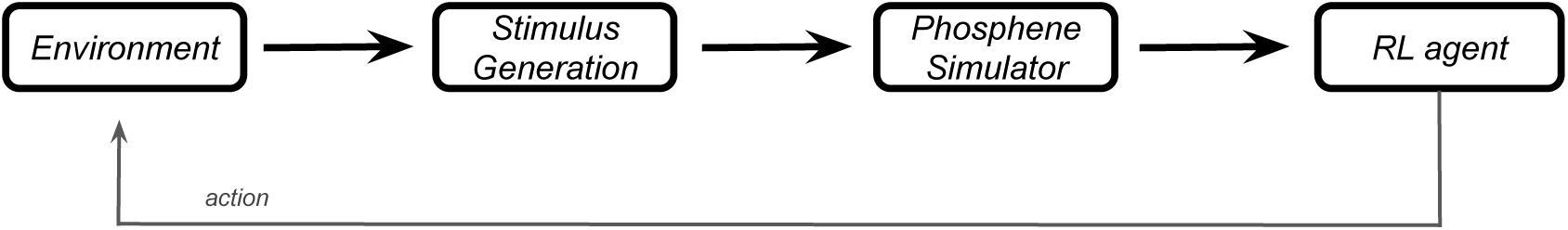
Pipeline for the task-oriented RL-based phosphene vision evaluation framework. First, the view of the virtual agent’s environment is captured to be provided for the stimulus generation process. A phosphene simulator transforms the output of the stimulus generation into phosphene vision. Based on this perception, the RL agent takes an action within the environment towards a task goal. The resulting movement changes the virtual agent’s view of the environment, allowing for another loop in the pipeline, which gradually converges on optimal decision making of the agent with the provided phosphene vision.

Detailed information about the elements of this evaluation framework can be found in Section 3.1, where possible choices for these elements are suggested within our TOPhos approach. Going beyond its role as an evaluation method, TOPhos showcases the RL-based evaluation framework’s additional capacity to optimize the phopshene generation process based on the task, opening up new possibilities in reaching ideal stimulation patterns for inducing phosphenes.

One advantage of this RL-based phosphene vision evaluation framework derives from the flexibility to choose the composition of each of its elements. It enables experimentation with any phosphene generation method in a range of environments and tasks, with the opportunity to explore their behavioral consequences before any testing with the actual neuroprostheses is possible. It provides a powerful test-bed for different phosphene generation methods and their potential effects

### 2.2 TOPhos: Task Optimized Phosphene Vision

TOPhos, our framework for task-dependent end-to-end optimization of phosphene vision through reinforcement learning, is composed of four architectural components: an encoder, a phosphene simulator, an RL agent and a decoder (Fig. 2), where the encoder handles the stimulus generation with help from the additional decoder component. The encoder takes frames from the environment scene as input and provides a stimulation pattern for inducing phosphenes, which corresponds to the electrical stimulation provided to a patient’s retina or visual cortex. The phosphene simulator in turn transforms this encoded stimulation into a representation of the phosphene percept called simulated phosphene vision which mimics the perception created in the patient’s mind as a result of the electrical stimulation. The actor-critic RL agent takes this phosphene representation as the sensory input of the current environment state, and outputs action probabilities and state value estimates to base its action on. After taking an action, the agent receives feedback from the environment in the form of a reward, which contributes an RL loss term to the optimization process. The simulated phosphene vision is also passed to the decoder, which interprets the phosphenes to reconstruct the original input frame. The reconstruction loss that emerges from the comparison of the original frame and the reconstructed frame contribute the second loss term during optimization. The combined reconstruction and RL losses direct the learning process of the agent simultaneously towards both improved task performance and interpretable phosphene patterns. The result is a task-specific end-to-end optimization of the phosphene vision. We provide further details on the individual components of TOPhos in the following subsections.^1^

**Figure 2:**
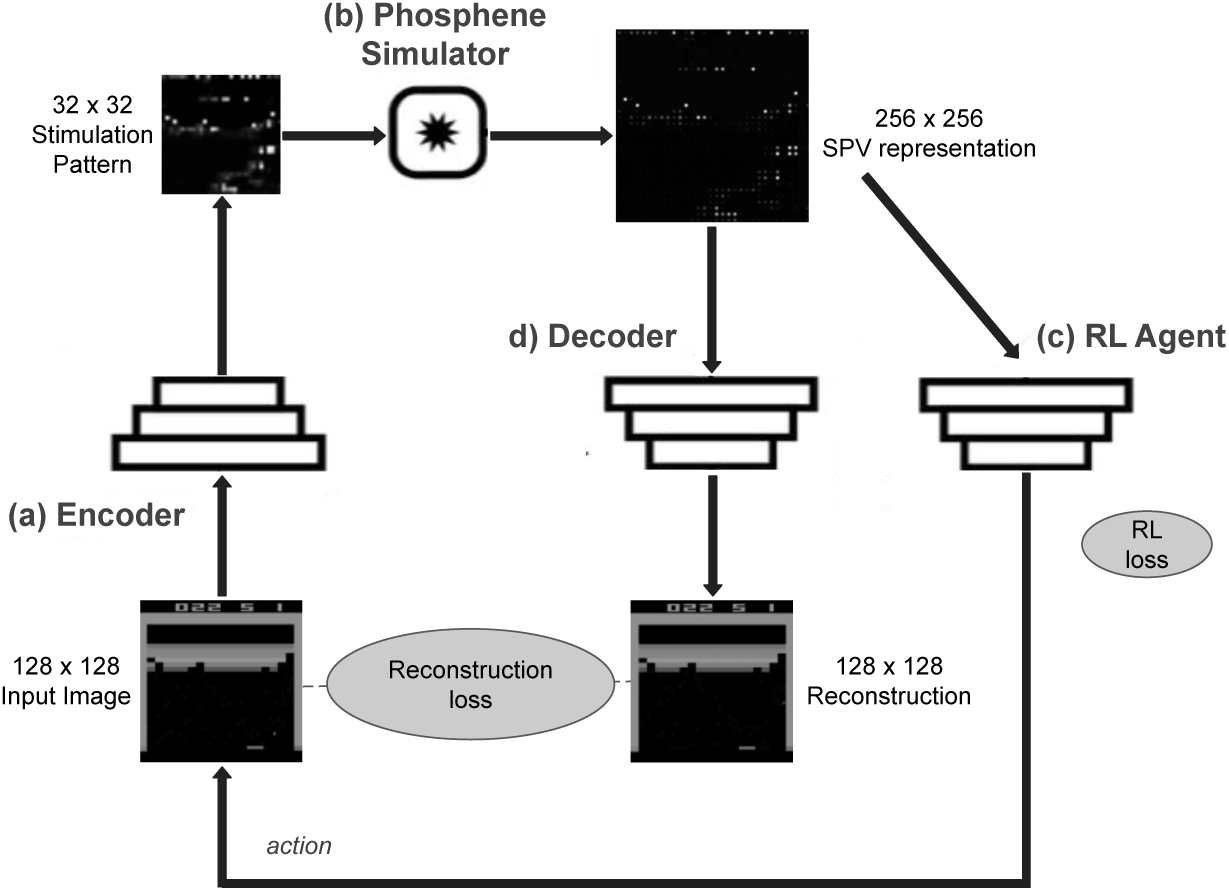
Architectural components of task-dependent end-to-end optimization of phosphene vision with processed Atari frames. (a) An encoder converts input frames into a 32 *×* 32 stimulation pattern. (b) A phosphene simulator captures the perceptual effect of the stimulation pattern and converts it into a simulated phosphene vision (SPV) representation of size 256 *×* 256. (c) The RL agent receives the SPV representation as a sensory input and uses it to compute actions in the environment. The actions produce an RL loss used to optimize the RL agent, and optionally the stimulus generation through the encoder, for the task at hand. (d) A decoder attempts to reconstruct the input frames from the SPV representation. The output of the decoder can be used to compute a reconstruction loss, which can be combined with the RL loss to drive the whole pipeline towards successful task performance with optimally generated phosphenes.

#### 2.2.1 Encoder for Stimulus Generation

The encoder is based on the model proposed by [17] for end-to-end optimization of phosphene vision. It is a convolutional deep neural network architecture (Fig. 3) including residual blocks. All convolutional layers use a kernel size of 3, with strides 1 and padding 1. We found RL performance to be more stable when using a Swish activation function after every convolution layer instead of batch normalization and leaky ReLU as in the original implementation [17]. The last convolutional layer is followed by a tanh activation, after which the outputs are scaled to the range [0, 1]. The encoder takes a frame of 128 *×* 128 as input and converts it into a spatial stimulation pattern of size 32 *×* 32. In a neuroprosthesis, this 32 *×* 32 grid corresponds to the implanted electrode array, where each electrode injects an electric charge proportional to the activation values at the output feature map of the encoder.

**Figure 3:**
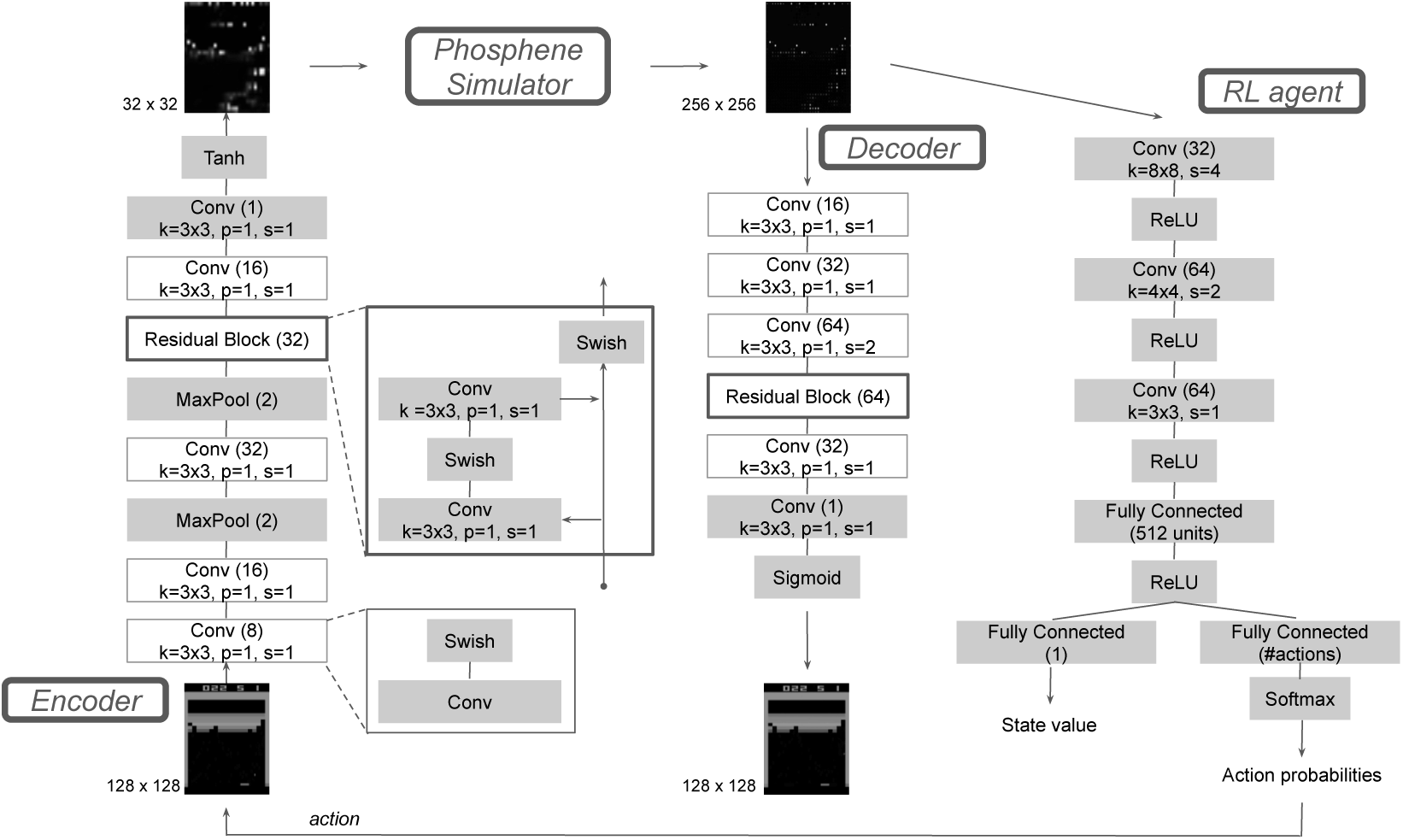
TOPhos: Architectural details of the neural networks used in task-dependent end-to-end optimization of phosphene vision. Grey boxes represent individual layers, white boxes are layer blocks.

#### 2.2.2 Phosphene Simulator

The phosphene simulator captures the perceptual effect of cortical stimulation in two steps [17]. First, the 32 *×* 32 stimulation pattern is upsampled to a 256 *×* 256 array by homogeneously inserting zeros between elements. Second, the resulting image is convolved with a Gaussian kernel, providing the approximated perceptual effect of electrical point stimulation. The increased resolution is chosen to obtain a simulated visual field large enough to contain phosphenes with characteristic Gaussian shapes. The phosphene simulator has no trainable parameters. Details about other parameters can be found in Appendix A.

#### 2.2.3 Reinforcement Learning Agent

The RL agent is an actor-critic model [24] consisting of a single-branch deep neural network with two fully-connected output layers (Fig. 3). The first output layer represents the actor and contains a neuron for each possible action. By applying a softmax on this output, the actor implements a policy through sampling from the resulting probability distribution over the action space. The second output layer represents the critic and consists of a single neuron generating the state value estimate. Overall, the RL agent takes a 256 *×* 256 *×* 4 representation of the environment’s state as input. Stacking the last four input frames is a common way to obtain richer spatiotemporal features and facilitate, e.g., motion estimation by the agent [25].

#### 2.2.4 Phosphene Decoder

The decoder is a convolutional residual network architecture inspired by previous work on end-to-end optimization of phosphene vision [17]. All convolutional layers use a kernel size of 3, with strides 1 and padding 1, except the third convolutional layer which uses a stride of 2. As in the encoder, all hidden convolutional layers are followed by a Swish activation function. The last convolutional layer uses a sigmoid activation to obtain normalized pixel values. The decoder receives a simulated phosphene representation of size 256 *×* 256 and returns a reconstruction of size 128 *×* 128 of the original input frame.

#### 2.2.5 Optimization Procedure

During training, the encoder, decoder and RL agent are all trained simultaneously in an end-to-end fashion. The learning algorithm is based on the PPO method [26], where multiple epochs of stochastic gradient descent are performed in each policy update for a batch of data collected in multiple parallel environments simultaneously. The PPO loss is composed of the actor loss 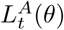, the critic loss 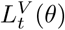 and the entropy loss 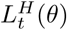. Together, they form what we call RL loss in this paper. In addition to this standard RL loss we introduce a reconstruction loss term 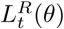, which arises from the mean squared error between the original frames and the reconstructed frames (Fig. 2). Combining these loss terms gives the objective as

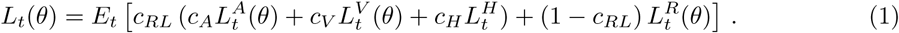

The coefficients *c*_*A*_, *c*_*V*_, *c*_*H*_ weigh each loss term in the PPO loss, whereas the coefficient *c*_*RL*_ trades off the overall RL loss against the reconstruction loss. *E*_*t*_ denotes the empirical mean over a set of samples. Details about the hyperparameters can be found in Appendix B.

## 3 Experiments & Results

To evaluate the performance of different stimulus generation approaches in our RL-based phosphene framework, we selected three simulated virtual environments from the Atari 2600 benchmark, namely Breakout, Seaquest, and Riverraid. These environments all require the agent mobility and adaptation to dynamically changing surroundings, which emulates scenarios encountered by users of a neuroprosthesis. However, the environments vary in difficulty and solution strategy, thus providing a way to test the task-optimality of our encoder. Game difficulty in this context is defined by the performance of DQN [24] relative to human performance, which is 25%, 57%, and 1317% in Seaquest, Riverraid, and Breakout.

Breakout was chosen as an environment where a fixed feature extractor like edge detection is expected to perform well because the success in gameplay prominently depends on capturing the view of a small ball that only encompasses a few pixels. The goal of the agent is simply to move a paddle horizontally to bounce the ball without dropping, so it can continue to break the bricks above. Seaquest is one of the most difficult games in the Atari benchmark in terms of achieving superhuman performance as it requires management of multiple goals, moving the agent in four cardinal directions while learning to collect, avoid or shoot various game objects. Riverraid follows it in this sense closely and is similar to navigating across a corridor that flows vertically while avoiding or aiming for objects, as in everyday situations of real agents.

The raw Atari frames of 210 *×* 160 pixels in RGB color were resized to 84 *×* 84 and converted to grayscale, before being used as input to the phosphene generation pipeline. All agents were trained until convergence and reward scores are reported based on averages over five runs with random seed initializations. Game plays were evaluated quantitatively based on performance of the trained agents, as well qualitatively by visual inspection of the simulated phosphene vision.

### 3.1 Task-dependent End-to-end Optimization of Phosphene Vision via Reinforcement Learning

In our main experiment, an RL agent was trained using the TOPhos architecture in Fig. 2. The sensory input to the RL agent is given by simulated phosphene vision, continuously adapted through the RL-based framework. The encoder, decoder, and RL agent are trained in an end-to-end manner by simultaneous optimization of task performance and reconstruction error. Fig. 4 provides some example phosphenes and reconstructions. ^2^ The weight of the RL loss relative to the reconstruction loss in Eq. 1 was determined via hyperparameter optimization as *c*_*RL*_ = 0.3. Giving more weight to the RL loss was found to suppress overall phosphene intensity. Turning off the reconstruction loss entirely (*c*_*RL*_ = 1) led to vanishing gradients, which is in line with previous interpretations of auxiliary losses acting as regularizers [21].

**Figure 4:**
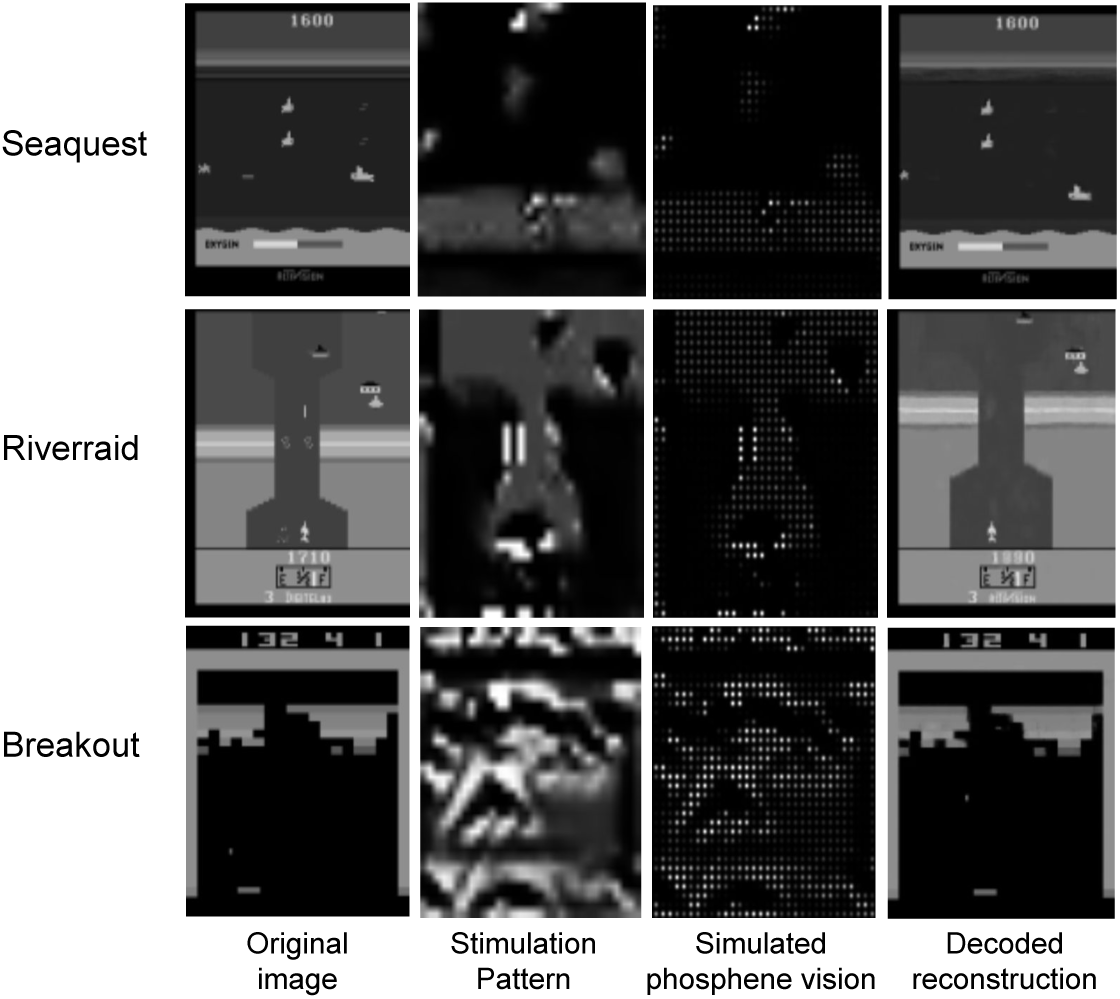
Example frames from each game and their transformation to phosphenes through training with TOPhos optimization.

### 3.2 RL Agent with Phosphene Vision Generated by Edge Detection

In this experiment, the RL agent acts on simulated phosphene vision generated by an edge detector. The mechanism for stimulus generation is fixed: Game frames are processed by an edge filter before being passed through the phosphene simulator and fed to the RL agent. We employ the OpenCV Canny edge detection implementation with a low threshold of 20 and high threshold of 40 [27].

Prosthetic vision based on edge-detection has shown promising results in navigation and object recognition scenarios [28, 29]. This experiment demonstrates the performance of an edge-based phosphene vision in our RL-based evaluation framework and investigates the value of static edge detection when evaluated in a task-dependent context.

### 3.3 Baseline Comparisons

Using the RL-based evaluation framework, we compared the performance of agents acting on phosphene vision against agents with other forms of vision as explained below. Training performance of agents with phosphene vision against these baselines can be seen in Fig. 5, showing final average scores and final maximum scores reached by the agents.

**Figure 5:**
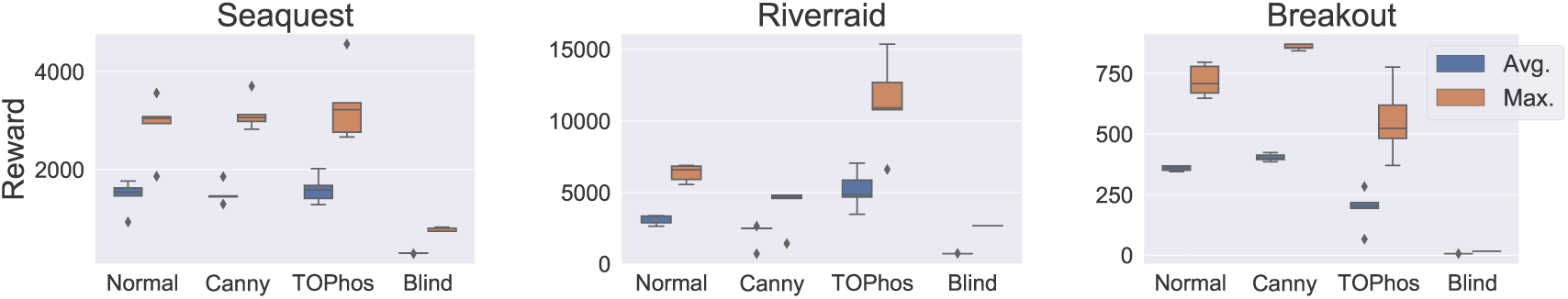
Distribution of average and maximum reward values obtained at the end of training on Seaquest, Riverraid and Breakout (games ordered from most to least difficult). The agent with end-to-end optimized phosphene vision (TOPhos) is compared against an agent receiving original input frames (Normal), no input frames (Blind) and phosphenes based on edge-detection (Canny). TOPhos and Canny show complementary performance: They are either on par or work best where the other is lacking.

#### 3.3.1 RL Agent with Normal Vision

The first baseline considers the performance of an RL agent with sensory input coming as close to normal vision as possible. This agent receives the resized grayscale game frames as input, without further pre-processing or phosphene simulation. Note that normal vision in this context is not a proxy for peak performance.

#### 3.3.2 Blind RL Agent

This experiment provides a baseline performance of an RL agent taking actions based on empty input, as a blind agent without access to other sensory modalities would do. Note that while this agent is expected to provide a lower bound on performance, a blind agent is able to learn based on reward signals and thus could in principle outperform an agent acting randomly. The blind agent provides a baseline for the performance gain to be expected from prosthetic vision.

### 3.4 Analysis of Baseline Comparisons

Fig. 5 shows that both types of phosphene vision (TOPhos and Canny) provide significant improvement in performance against the blind agent in all games, based on two-tailed t-tests (*p <* 0.05, *N* = 10). Agents with phosphene vision can reach a level of performance comparable to the agent with normal vision. Phosphene vision even appears advantageous over normal vision in case of Riverraid (using TOPhos features) and Breakout (using edge features), suggesting a benefit of using more expressive features and reduced input complexity.

On the game Seaquest, the means between both TOPhos and Canny-based phosphene vision do not differ significantly according to a t-test. A significant improvement is observed with TOPhos on Riverraid, and with Canny on Breakout. The advantage of Canny-based phosphene vision in the more simplistic game Breakout is expected given that the agent only needs to detect and minimize its distance to the ball, for which an edge filter provides reliable and sufficient features. Using edge features can be seen as applying the right inductive bias during representation shaping [21]. The same inductive bias, that is, reliance on edge features, appears insufficient in the more challenging game Riverraid. The indiscriminate application of an edge filter is unable to remove redundant features that are task-irrelevant. Examples can be seen in Fig. 6, where the static borders of the game frame or the scoreboard are encoded with phosphenes in Canny, while they were learned to be ignored in TOPhos, thus improving sparsity.

**Figure 6:**
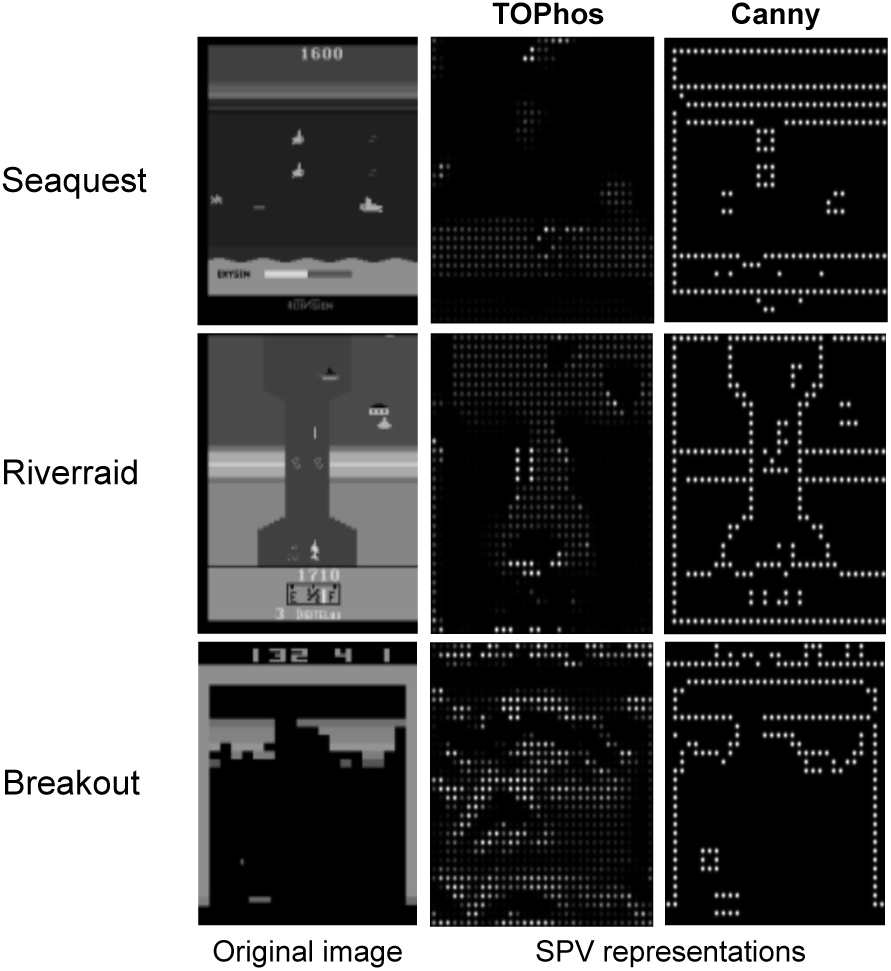
Example input frames and respective simulated phosphene vision from the E2E-optimized TOPhos model and the canny edge detection.

The end-to-end optimized phosphene vision performs better than fixed edge-based phosphene vision in Riverraid and on-par in the similarly complex game Seaquest. This observation suggests that by adapting the stimulus generation towards task demands, the end-to-end optimization provides performance gains when the task is visually complex and requires handling multiple goals simultaneously. When the success in the game depends largely on accurately capturing a single pixel such as the ball in Breakout, a direct mechanistic stimulus generation may be more effective, whereas multiple mobile task-relevant details are better emphasized and distinguished with an adaptive end-to-end optimized approach.

### 3.5 Studying the Components of the Optimization Objective

We studied the contribution of reconstruction and RL losses for optimization of the phosphene vision via end-to-end training. Specifically, we isolated the effect of optimizing towards the task demands through the RL loss, and towards accurate phosphene decoding through the image reconstruction loss. As performance metric we used the final reward averaged over five runs on the Breakout task.

#### 3.5.1 Effect of RL Loss on Optimization of Phosphenes

To highlight the contribution of task-dependent optimization via an RL loss, we removed the RL agent from the TOPhos architecture. The model then consists of the encoder for stimulus generation, the parameter-free phosphene simulator, and the decoder for input reconstruction. The encoder and decoder were trained for 20 epochs using only the reconstruction loss from decoded phosphenes. The input consisted of 4500 original game frames in batches of size 20. The frames were sampled from the gameplay of an independent agent pre-trained with normal vision.

After training, the encoder was fixed, the decoder was removed, and an RL agent was trained to solve the task given phosphenes from the pre-trained encoder. The resulting performance (Fig. 7) informs about the quality of phosphenes when the encoder is trained without a task-focus.

**Figure 7:**
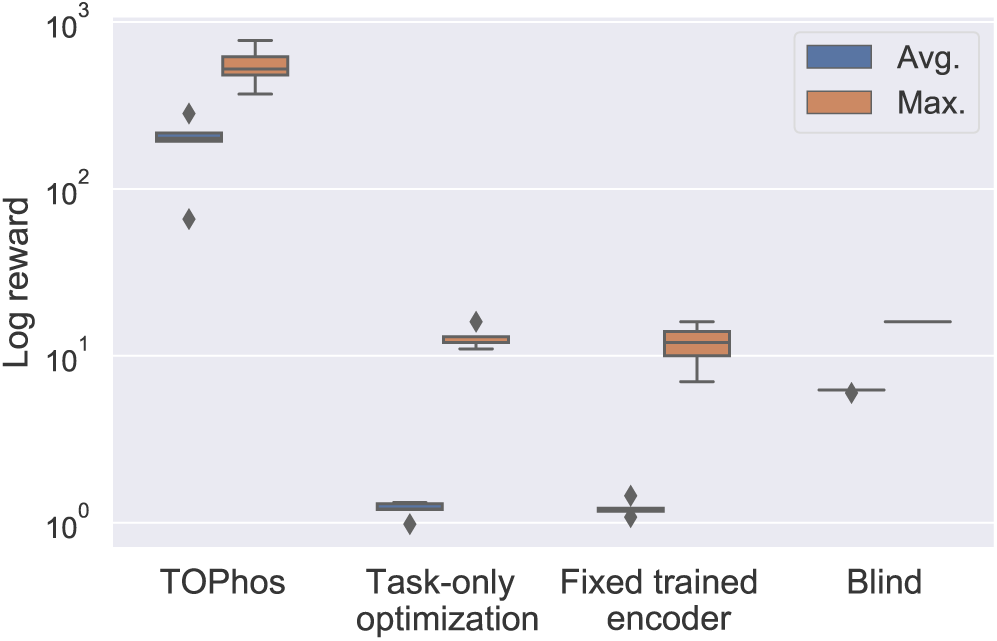
Distribution of average and maximum logarithmic reward values obtained at the end of training on Breakout. The agent with end-to-end optimized phosphene vision (TOPhos) is compared against the blind agent (Blind) and two variants of TOPhos: ‘Task-only optimization’ denotes training without the decoder and ‘Fixed trained encoder’ trains an RL agent with a fixed encoder trained beforehand without the RL loss.

Despite successful minimization of the reconstruction loss, the agent falls short of the performance of the agent whose encoder is trained with a task focus. The state representation with the given phosphenes is not informative enough for the agent to learn the game. Fig. 8 illustrates the phosphene patterns generated by an encoder trained only on the reconstruction loss, without having the RL agent in the loop. The phosphenes are less sparse than those learnt through task-dependent end-to-end optimization (see Fig. 4 for a comparison). Note that we do not include an explicit sparsity regularizer in our objective function as in previous work [17]. This observation suggests that optimization for a task may lead to the benefit of sparser representations in addition to providing more informative task-relevant features to the RL agent.

**Figure 8:**
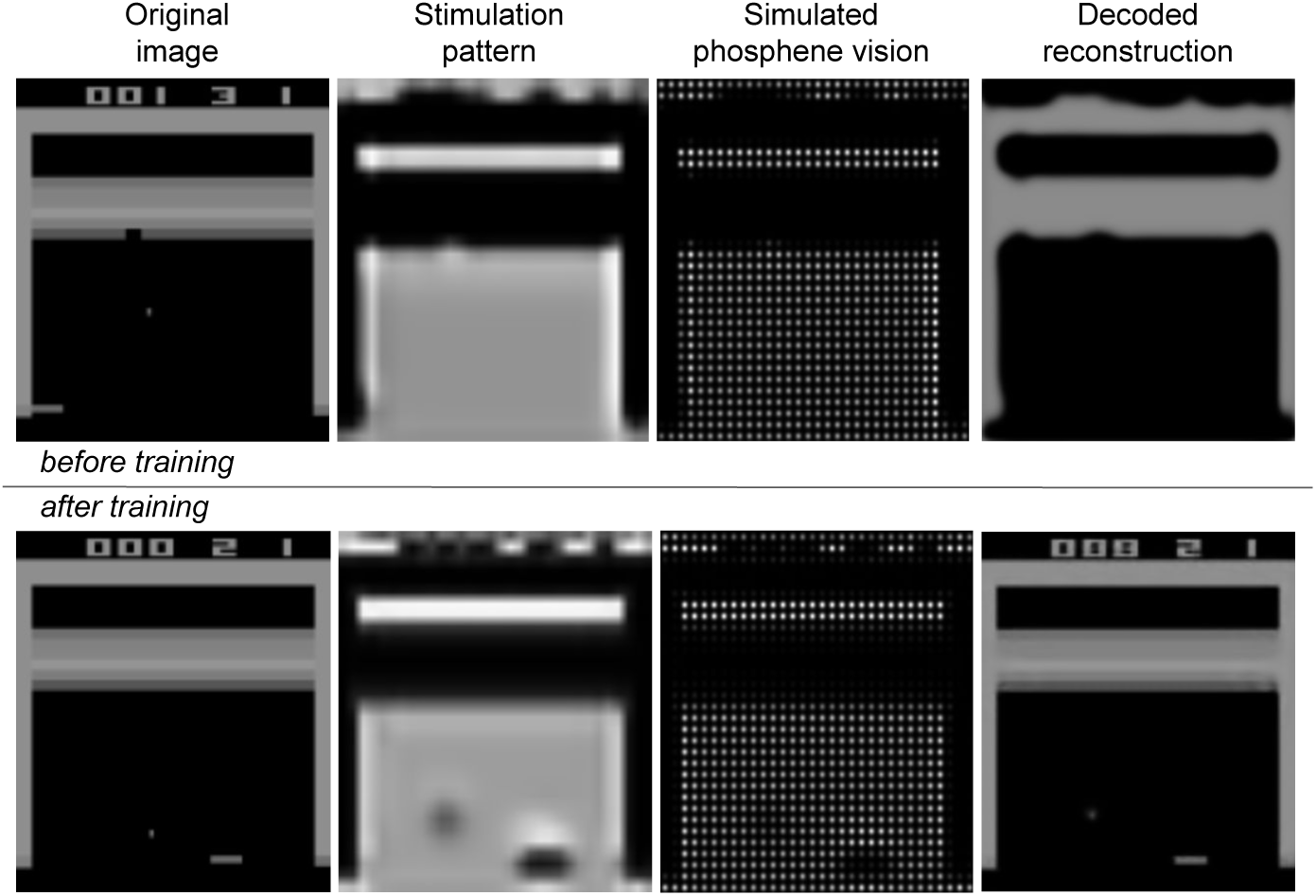
Phosphene patterns and reconstructions before and after training the encoder using reconstruction loss only, without a task focus. The resulting stimulation and phosphene patterns lack sparsity.

#### 3.5.2 Effect of Reconstruction Loss on Optimization of Phosphenes

This experiment investigated whether optimizing only for the task is enough for meaningful phosphene vision. The decoder was removed from the TOPhos architecture, hence no reconstruction loss was available. The model then consists of the encoder for stimulus generation, the parameter-free phosphene simulator, and the RL agent. The encoder and RL agent are trained together with the objective to optimize the RL loss only.

As seen in Fig. 7, the performance of this agent is also worse than the fully end-to-end optimized RL agent, demonstrating the benefit of including an auxiliary reconstruction loss in the optimization. In fact, training without reconstruction loss was found to result in vanishing gradients and suppressed phosphenes. Taken together, these findings justify the use of both reconstruction loss and task-dependent RL loss in our end-to-end optimization framework for phosphene vision.

### 3.6 Learning Task-Specific Features

To verify that the stimulation patterns learned through the RL-based end-to-end optimization are indeed task-specific, we used the dimensionality reduction technique t-SNE [30] to visualize the clusters emerging when generating patterns from frame inputs of a single game using encoders trained on different games. From the end-to-end TOPhos models trained as described in Section 3.1, we took the encoders from best performing agents in Seaquest, Riverraid, and Breakout. Each of these task-specific encoders was applied on sample frames from all three games. The stimulation patterns resulting from inputs of a given game were visualized using t-SNE (see Fig. 9). For this, dimensions were first reduced to 50 using PCA, then t-SNE was applied with a perplexity value of 20 to arrive at a representation of stimulation patterns in 2-dimensional space. For example, the first panel in Fig. 9 represents the patterns obtained when feeding Seaquest frames to the three encoders trained on Breakout, Riverraid, and Seaquest, respectively.

**Figure 9:**
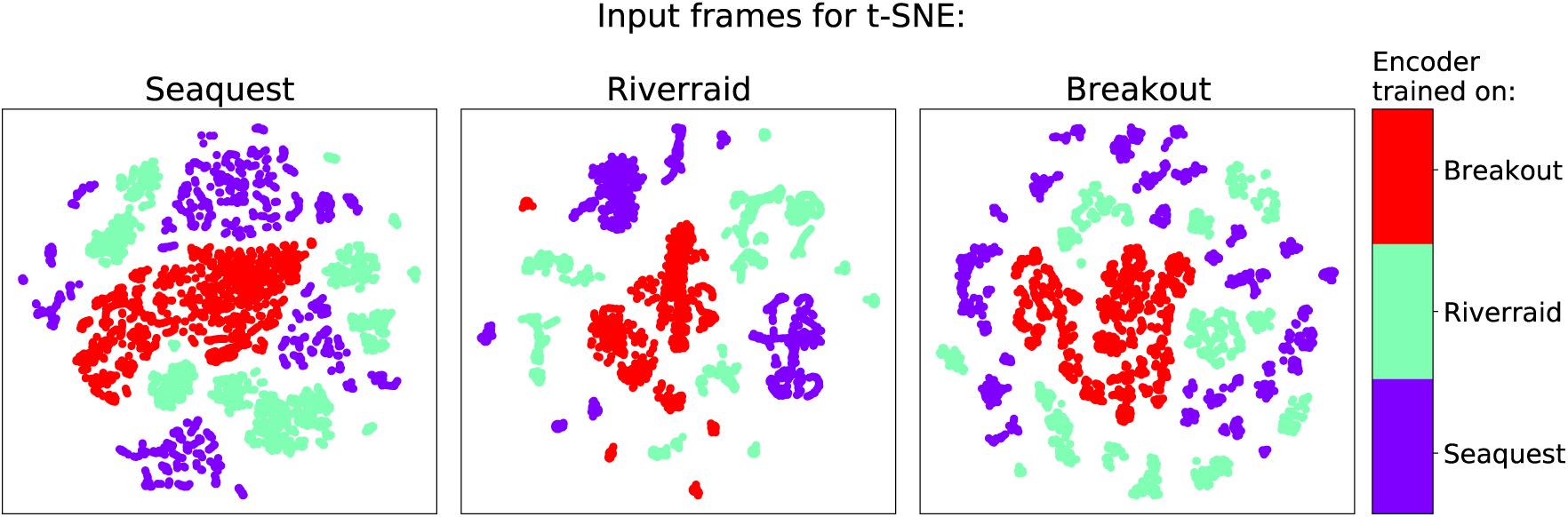
T-SNE clusters of stimulation patterns obtained from encoders trained on different games (colors in legend) when applied on input frames of a given game (from left to right: Seaquest, Riverraid, Breakout).

If the encoders were task agnostic, then there would be no clusters in t-SNE and t-SNE representations of different colors would overlap. Instead, we observe distinct clusters for each encoder. It confirms that the stimulation patterns learned as a result of the end-to-end RL-based optimization process are indeed shaped towards the task requirements.

Finally, we qualitatively compared stimulation patterns generated from an encoder when receiving frames from the game it was trained on versus frames from a different game. Identifying which environment the scene belongs to is easier when using the encoder trained on that environment. The stimulation pattern becomes less informative when applying the encoder trained on another environment (for an illustration see Appendix C).

Overall, these findings confirm the learning of task-specific features by the encoders as a result of the end-to-end optimization process with a task-oriented RL framework.

## 4 Discussion

In this work we propose TOPhos, an optimization framework for prosthetic vision where the encoder for stimulus generation is adapted to the requirements of a given task. To this end, the encoder is trained end-to-end with a reinforcement learning agent in the loop. Our results demonstrate that the resulting task-dependent encoder performs on-par with fully sighted agents and can offer benefits compared to fixed stimulus generation methods such as edge detectors. In addition, we show that an auxiliary loss term using image reconstruction from decoded phosphenes is an essential ingredient to sparse and informative phosphene vision.

The proposed framework for phosphene optimization in an RL setting is well suited to flexibly evaluate different representation learning methods in simulation before moving to behavioral studies in humans. As a first approximation, one can take the performance of a virtual agent as proxy for real agents in naturalistic environments when receiving phosphene vision either via a static preprocessor such as edge detection or semantic segmentation, or via a task-adaptive feature extractor as proposed here. Our RL-based evaluation revealed that end-to-end optimized TOPhos and fixed Canny encoders can be complementary; the proposed optimization framework is a valuable test bed to explore combinations of both approaches in subsequent studies.

When comparing end-to-end optimized phosphene vision against fixed edge-based phosphene vision, the TOPhos approach matches or surpasses Canny performance in the two difficult games tested, yet underperforms in the simpler game Breakout. It suggests that TOPhos is suited for settings where the task is visually complex and requires distinguishing multiple object types and behavioral patterns. Breakout differs from the other games because high scores can be reached by meeting a single goal: Shifting a paddle horizontally to prevent a ball from dropping below the game window. For this task, an edge detector that reliably picks up the location of ball and paddle provides a reasonable inductive bias for representation learning. In the TOPhos framework, the absence of a single-pixel ball has little impact on the reconstruction error, which sometimes leads to suboptimal training results. In contrast to Breakout, both Riverraid and Seaquest require juggling multiple goals at the same time, and distinguishing which objects in the scene to avoid as enemies or approach for additional reward or survival. In this scenario, the indiscriminate nature of the edge filter may make the task relevance of game objects harder to interpret and is sometimes unable to eliminate non-informative static visual elements in the scene. On the other hand, the encoder learned in the end-to-end TOPhos approach appears to accentuate moving objects and allow sparsification of the phosphene representation even without an explicit sparsity loss. This sparsity benefit is crucial for reducing the energy consumption of a neuroprosthesis, as well as maintaining stimulation levels that are continually sustainable by the neural tissue. Another possible disadvantage of an edge-based feature extractor is its inflexibility to changes in overall contrast or lighting, which may require an adaptation of the threshold parameters. While a definite conclusion cannot be drawn from the environments considered here, our study indicates that an end-to-end task-dependent optimization scheme can be beneficial in scenes with higher complexity, object variability, and variance of image statistics, for which a parametrized encoder can adapt during training.

An exciting avenue for future research will be to incorporate bio-inspired models in the phosphene generation pipeline, and compare the simulated phosphene vision with neurological data on induction and perception of phosphenes in behavioral studies.

Though Atari games may seem distant from typical situations encountered by the visually impaired, Riverraid and Seaquest share characteristics that make them relevant to the use case of prosthetic vision, namely visually guided navigation, object manipulation, and goal-directed behavior in complex and dynamic environments. While already evident here, we expect that the advantage of end-to-end RL-based phosphene optimization will become fully apparent when evaluated in naturalistic environments. The present study lays the ground for this future work and contributes to the optimal control of stimulation patterns in visual neuroprosthetics.

## Acknowledgements

This work has received funding from the European Union’s Horizon 2020 research and innovation programme under grant agreement No 899287.

## A Phosphene Simulator Hyperparameters

**Table 1:** Parameters used for the phosphene simulators.

| Parameter | Value |
| --- | --- |
| Phosphene density | 32 |
| Gaussian kernel size | 11 |
| Gaussian kernel $\sigma$ | 1.5 |
| Intensity | 15 |
| Upsampling scale factor | 8 |

## B TOPhos Hyperparameters

**Table 2:**
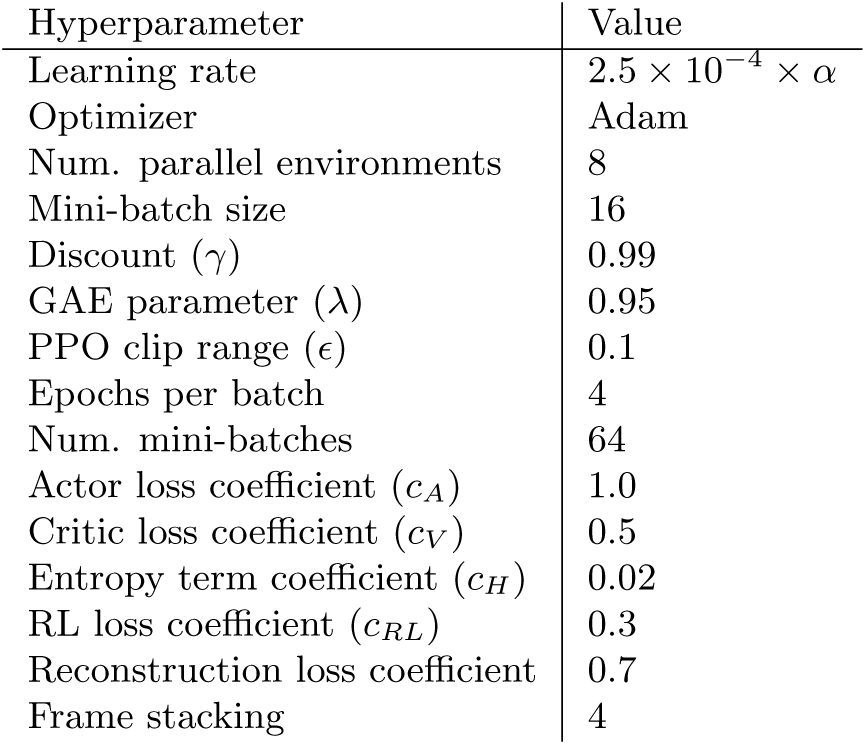
Hyperparameters used in training the TOPhos agents, where *α* is linearly annealed from 1 to 6 *×* 10^*−*5^ over 15000 iterations.

| Hyperparameter | Value |
| --- | --- |
| Learning rate | $2.5 \times 10^{-4} \times \alpha$ |
| Optimizer | Adam |
| Num. parallel environments | 8 |
| Mini-batch size | 16 |
| Discount ($\gamma$) | 0.99 |
| GAE parameter ($\lambda$) | 0.95 |
| PPO clip range ($\epsilon$) | 0.1 |
| Epochs per batch | 4 |
| Num. mini-batches | 64 |
| Actor loss coefficient ($c_A$) | 1.0 |
| Critic loss coefficient ($c_V$) | 0.5 |
| Entropy term coefficient ($c_H$) | 0.02 |
| RL loss coefficient ($c_{RL}$) | 0.3 |
| Reconstruction loss coefficient | 0.7 |
| Frame stacking | 4 |

## C Stimulation Patterns on Different Game Frames

**Figure 10:**
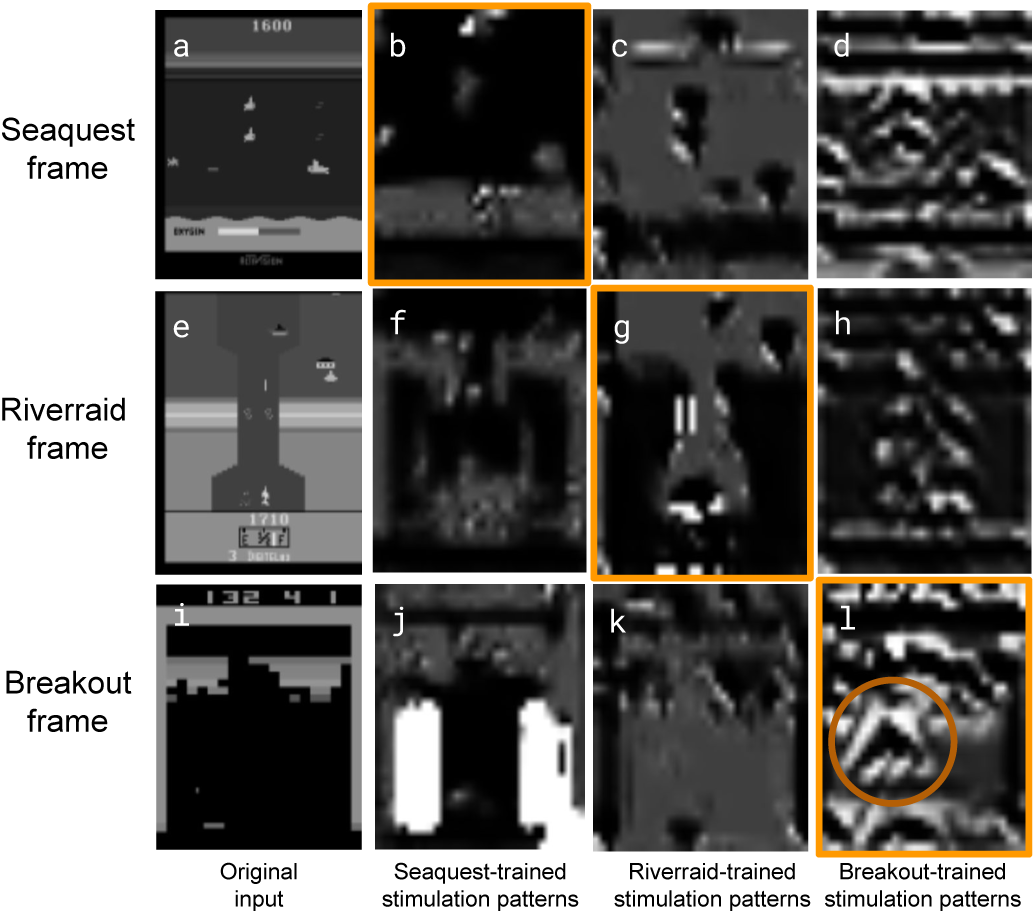
Example frames from a given game (first column) and respective phosphene representations produced using encoders trained on different games (from left to right: Seaquest, Riverraid, and Breakout). Matching games of input frames and encoder were highlighted in orange, where phosphene representations are more informative due to the task-adaptive optimization process. For example, for a Riverraid frame (e), only the matching encoder’s stimulation patterns (g) remind of the original input, enabling the detection of the vertical game flow since only the areas and objects where the agent is allowed to step on were encoded with phospenes. The Seaquest-trained encoder’s application on a Breakout frame (j) results in large and strongly activated regions which cause a distraction from relevant game objects. On the other hand, the Breakout-trained encoder learns to accentuate the movement of the ball by encoding a triangular arrow-like shape around it (highlighted with a circle in l), where the upper corner of the triangle points towards the desired direction of motion. This task-specific encoding becomes less informative when applied on frames from the other games (d, h). On Seaquest frames, the Seaquest trained encoder (b) displays a much sparser representation of the input compared to other encoders (c, d), thus conserving energy and improving interpretability.

## Footnotes

1 Code is available at https://github.com/burcukoglu/TOPhos.git

2 Videos of the performance of trained agents and their phosphene vision transformation can be seen in https://youtube.com/playlist?list=PLOsc1mo6nsPeBdfEvjSKL6_UWb-M2unJH

